# Histone H1 regulates non-coding RNA turnover on chromatin in a m6A-dependent manner

**DOI:** 10.1101/2021.10.12.464039

**Authors:** José Miguel Fernández-Justel, Cristina Santa-María, Alberto Ferrera-Lagoa, Mónica Salinas-Pena, Magdalena M. Maslon, Albert Jordan, Javier F. Cáceres, María Gómez

## Abstract

Linker histones are highly abundant chromatin-associated proteins with well-established structural roles in chromatin and as general transcriptional repressors. In addition, it has been long proposed that histone H1 exerts context-specific effects on gene expression. Here, we have identified a new function of histone H1 in chromatin structure and transcription using a range of genomic approaches. We show that histone H1-depleted cells accumulate nascent non-coding RNAs on chromatin, suggesting that histone H1 prevents non-coding RNA transcription and regulates non-coding transcript turnover on chromatin. Accumulated non-coding transcripts have reduced levels of m6A modification and cause replication-transcription conflicts. Accordingly, altering the m6A RNA methylation pathway rescues the replicative phenotype of H1 loss. This work unveils unexpected regulatory roles of histone H1 on non-coding RNA turnover and m6A deposition, highlighting the intimate relationship between chromatin conformation, RNA metabolism and DNA replication to maintain genome performance.

---

Linker histone H1 plays an essential role in the folding of nucleosome arrays into more compact chromatin structures. Importantly, growing evidence from the last decade supports the concept that histone H1 is a multifunctional protein that can block the binding of other proteins to chromatin and also act as a recruitment platform for activators or repressors thus fine-tuning chromatin function (1, 2). There are seven genes coding for somatic linker histone H1 subtypes or variants in the mouse and human genomes, with an average of 0.5 to 1.3 H1 molecule per nucleosome depending on cell type (3). Disruption of one or two linker histone genes, initially performed to delineate subtype-specific functions, revealed that cells can maintain their total H1 content through compensatory up-regulation of the remaining H1 genes (4). However, inactivation of three subtypes leading to a 50% of the normal level of H1 resulted in embryonic lethality in mice, demonstrating that a correct stoichiometry of linker histone deposition on chromatin is essential for mammalian development (5). Embryonic stem cells (mESCs) derived from these triple-knockout embryos (H1-TKO) have a genomic average of one H1 molecule every four nucleosomes. These cells display limited changes in gene expression, yet they display de-repression of major satellite elements (5–7).

The distribution of histone H1 throughout the genome is not uniform. It has been shown that chromatin at active and poised gene promoters is characterized by reduced histone H1 levels, while inactive genes and heterochromatin are enriched in H1 (6, 8–9). In addition, H1 mediates the silencing of heterochromatic repetitive elements both by modulating their higher-order structure, but also by interacting with the histone methyltransferases responsible for the repressive methylation of these regions (10). The dual role of H1 is not limited to heterochromatin, since it also affects chromatin architecture by interacting directly with both transcriptional activators and/or repressors. Some examples include H1 binding to Cul4A ubiquitin ligase and the PAF1 elongation complexes that help to maintain active gene expression (11), its recruitment by the Msx1 factor to a regulatory element in the MyoD gene resulting in repressed muscle cell differentiation (12), or its interaction with p53 repressing its transcriptional activation effect (13).

We previously found that reductions on histone H1 content generated genome-wide alterations in their replication initiation patterns, as well as massive fork stalling and DNA damage due to replication-transcription conflicts (14). These findings raised the question of how limited alterations in gene expression upon H1 deficiency can be reconciled with widespread replication-transcription conflicts. To delineate novel functions of histone H1 on transcription regulation here we performed a detailed analysis of chromatin transcript abundances, RNA polymerase II (RNAPII) location and activity, and nascent RNA N-6-adenosine methylation (m6A) profiling in H1-TKO deficient mESCs (triple knock-out for the subtypes *H1c*, *H1d* and *H1e*). We found that reductions in histone H1 content resulted in the presence of thousands of cis-regulatory non-coding transcripts bound to chromatin. These ncRNAs were actively transcribed, anchored to chromatin through RNAPII complexes, and displayed reduced levels of m6A. Remarkably, knockdown of the m6A de-methylases ALKBH5 and FTO, and the m6A reader YTHDC1, rescues the replicative stress of H1-TKO cells. These results indicate that an appropriate histone H1 content is required to limit ncRNAs accumulation on chromatin, likely by reducing both RNAPII recruitment and also by facilitating co-transcriptional m6A deposition. Our findings reveal an unexpected role of histone H1 in regulating non-coding RNA turnover in chromatin and uncover a novel link between chromatin conformation and RNA post-transcriptional modifications, with important implications for understanding genome functionality.

## RESULTS

### Reductions in histone H1 content lead to the accumulation of non-coding transcripts on chromatin

A variety of models of H1-depletion in different systems showed limited transcriptional alterations, comprising both up and down-regulation of specific sets of genes (5, 7, 15–19). To investigate the mechanism by which H1 deficiency leads to transcription-dependent replicative stress we searched for chromatin-enriched RNAs in H1-TKO mESCs, using the CheRNA-seq approach (20). This technique preferentially detect RNAs bound to chromatin through RNA polymerase molecules, thus enabling the analysis of partially processed transcripts, as well as of structural RNAs (**Figure 1a**). The enrichment of chromatin associated transcripts in CheRNA preparations was monitored by checking the chromatin/nucleoplasm ratio for *Kcnq1ot1* and *Neat1*, two noncoding RNAs that associate to chromatin post-transcriptionally, relative to *Klf16* and *Nat8L*, two normally exported mRNAs (20) (**Figure 1b**). Inclusion of a spikein luciferase RNA allowed to assess potential changes in the overall amount of chromatin-bound RNA between preparations (**Supplementary Figure 1a**). Normalized reads from triplicate experiments from WT or H1-TKO mESCs were used to build a *de novo* transcriptome, and transcripts were classified in four classes regarding their genome location and coding potential: i) internal antisense RNAs (IAS), ii) long intergenic non-coding RNAs (lncRNAs), iii) promoter upstream transcripts (PROMPTs), and iv) coding RNAs (coding) (**Supplementary Figure 1b**). CheRNA-seq analyses revealed thousands of differentially expressed transcripts comprising the four categories (fold change >2 and adjusted p-value <0.01) (**Figure 1c**). Strikingly, all non-coding classes were upregulated in H1-TKO cells (**Figure 1d-e**; blue, upregulated in WT, and red, upregulated in TKO). In addition, intergenic reads not statistically included in the *de novo* transcriptome were also higher in H1-TKO cells (**Supplementary Figure 1c-d**). These findings indicate that an appropriate histone H1 content is required to prevent the accumulation of non-coding transcripts in chromatin that could potentially interfere with DNA replication.

**Figure 1.**
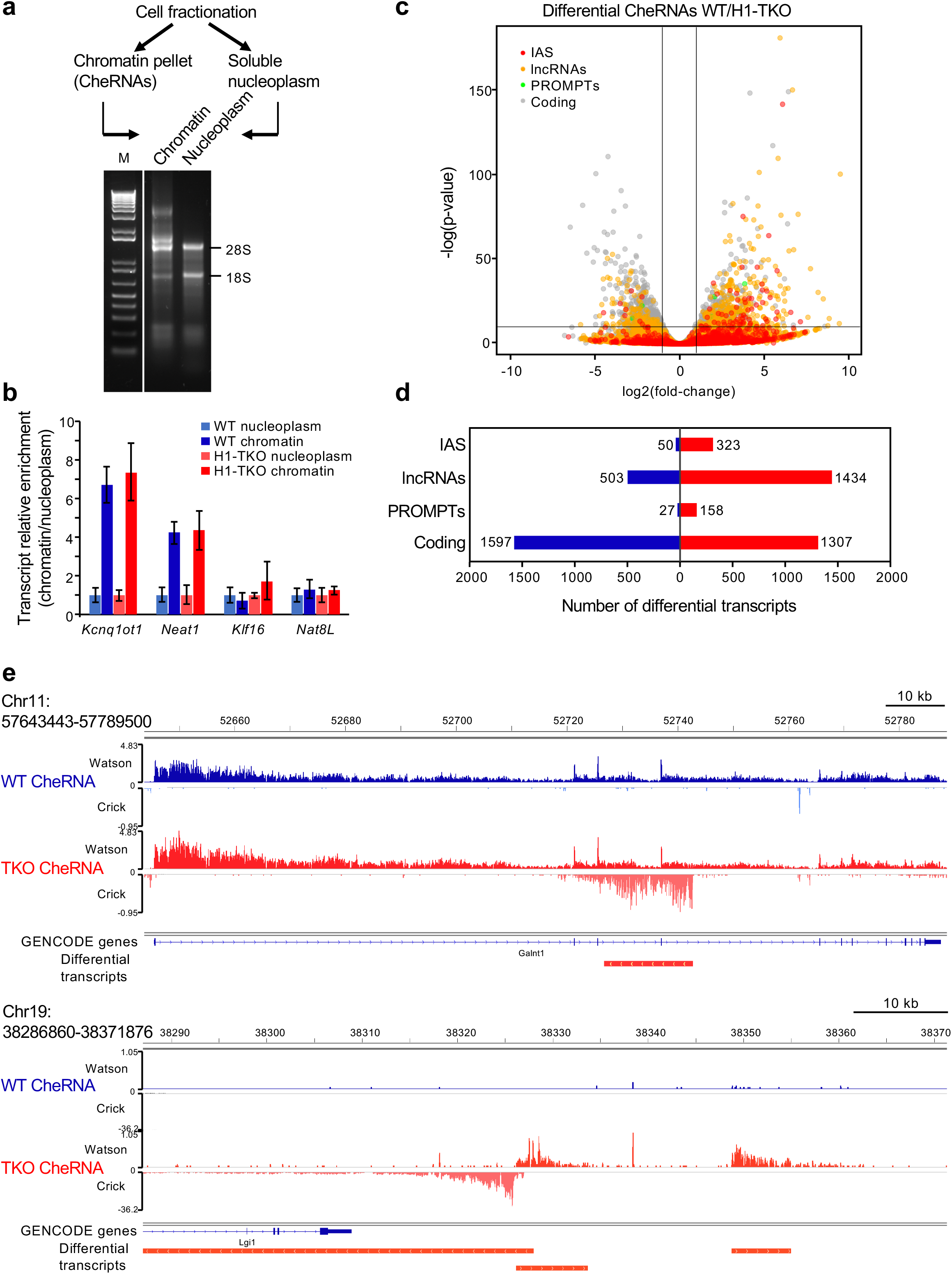
Histone H1-defficient cells accumulate non-coding transcripts on chromatin. **(a)** Schematics of CheRNA purification and representative example of chromatin and nucleoplasmic RNA fractions run in a non-denaturing 1% agarose gel. M, 1 kb ladder. **(b)** Ratio between chromatin and nucleoplasmic levels for the indicated RNAs: *Kncq1ot1* and *Neat1* lncRNAs associate to chromatin post-transcriptionally (Werner and Ruthenburg, 2015), and *Klf16* and *Nat8L* mRNAs do not. RT-qPCR data represent the mean and standard deviation of three biological replicates of WT and H1-TKO mES cells preparations (n=3). Blue, WT cells, red, H1-TKO. **(c)** Volcano plot showing the log2(fold change) and the −log(p-value) for the indicated CheRNA classes between WT and H1-TKO cells: internal antisense (IAS), lncRNAs, promoter upstream transcripts (PROMPTs) and coding RNAs. **(d)** Number of differentially expressed transcripts between WT and H1-TKO cells for each CheRNA category. Blue, overexpressed in WT; red, overexpressed in H1-TKO. **(e)** Representative IGV browser snapshots of transcripts specifically accumulated in the chromatin of H1-TKO cells. Upper panel, IAS for the *Galnt10* gene. Lower panel, several lncRNAs adjacent to the silent *Lgi1* gene.

### Accumulated non-coding transcripts are regulatory RNAs

Chromatin-associated non-coding RNAs frequently have *cis*-regulatory functions (21, 22). To evaluate whether the novel lncRNAs unveiled in mESCs with reduced histone H1 levels were indeed enhancing the activity of neighboring promoters, we first calculated the expression levels of coding genes relative to their distance to a differentially expressed lncRNA (20, 21) (**Figure 2a**). The analysis showed that the closer to a lncRNA, the higher the transcriptional activity of a gene. Consistent with this, a large fraction of up-regulated non-coding transcripts were generated from enhancer-like chromatin regions as previously defined (23) (**Figure 2b** and **Supplementary Figure 1e**), and displayed enrichment in H3K4me1 around their transcription start sites (TSS) (**Figure 2c**). Likewise, lncRNAs with differential enrichment in H1-TKO cells were preferentially located *in cis* of genes functionally involved in development and RNA polymerase II (RNAPII) transcription (**Figure 2d**). A deeper analysis of this last group revealed that it comprised key transcription factors involved in cell differentiation, including the pluripotency factors *Sox2*, *Nanog*, *Oct3/4* and *c-Myc*. In all cases, the lncRNAs TSS’s mapped at the superenhancer regions that regulate the transcription of these genes in embryonic stem cells (24–26) (**Figure 2e**). We subsequently examined the dbSUPER database and found that, out of the 231 annotated superenhancers in mESCs (27), 227 matched an assembled lncRNA. Thus, the chromatin-associated lncRNAs unmasked when histone H1 levels are reduced fulfill the requirements of regulatory RNAs. As the functional terms of the differential lncRNAs-neighbor genes were similar to the set of differentially expressed coding genes (**Supplementary Figure 1f**), we checked whether the accumulation of non-coding transcripts in H1-TKO chromatin was altering the expression of the proximal coding gene. We found that the foldchange between lncRNA expression and that of its neighbor coding gene was correlated for down-regulated, but not for up-regulated lncRNAs (**Figure 2f** and **Supplementary Figure 1g**). Examples of each scenario are shown in **Figure 2e**. A similar trend was detected for IAS and PROMPTs, despite their lower numbers (**Supplementary Figure 1h-i**). Collectively, these data suggest that histone H1 is a repressor of (silent) regulatory lncRNAs.

**Figure 2.**
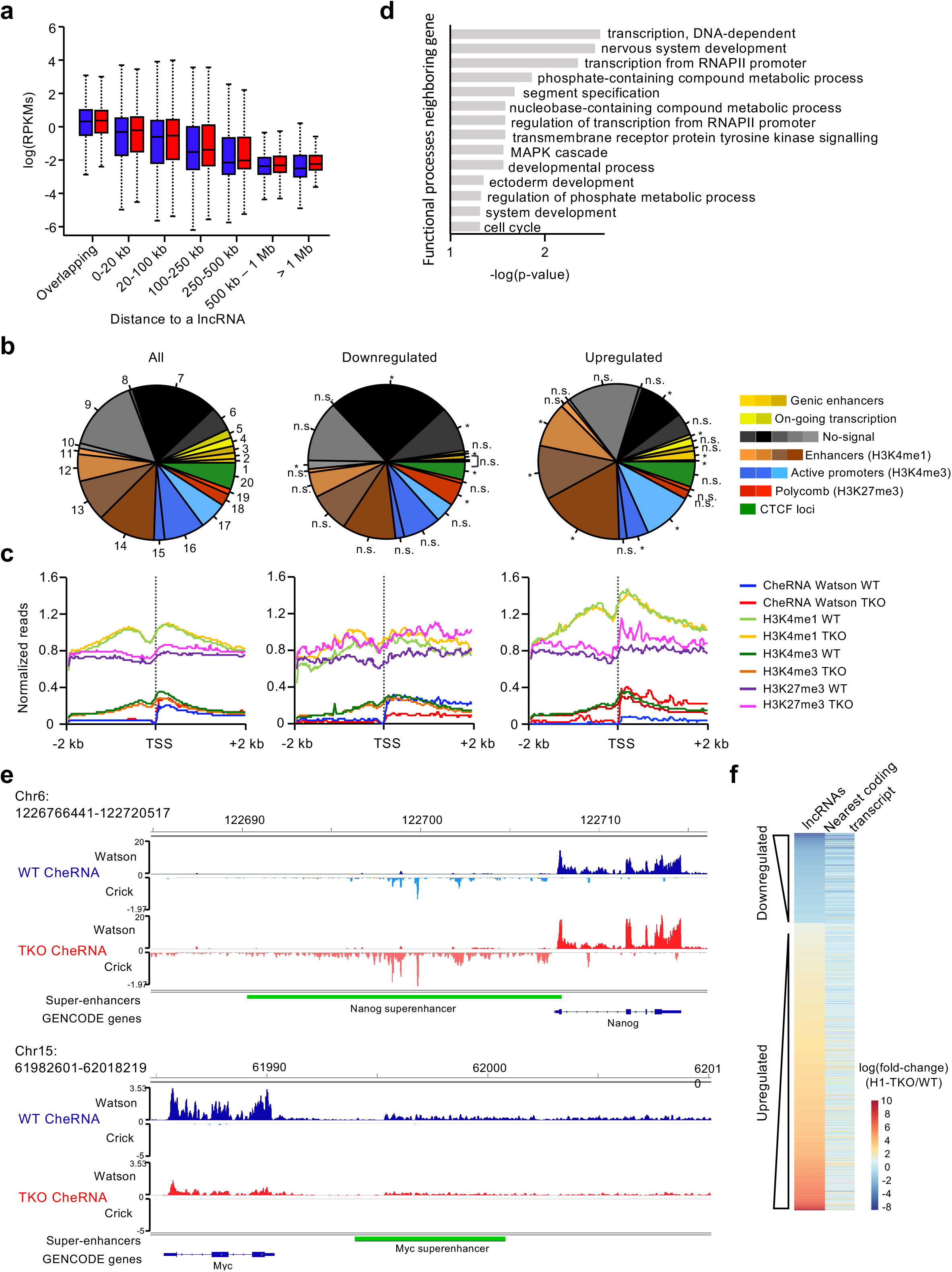
Accumulated non-coding transcripts are *cis*-regulatory RNAs. **(a)** Distribution of log(RPKMs) of coding genes located at the distances shown of a lncRNA. Blue, WT cells, red, H1-TKO cells. **(b)** Percentage of lncRNAs whose TSS matches each of the chromatin states shown. The percentage was compared with the expected one obtained from 100 random permutations of the differential transcripts, and the p-value was calculated. *p-value<0.01. Chromatin states are from Juan et al. (2016). Full description of the mESCs chromatin states is shown in Supplementary Figure 1e. **(c)** Profile of CheR-seq and ChIP-seq signal of the indicated epigenetic marks plotted in 4kb windows surrounding the TSS of the lncRNA’s categories shown. H3K4me1 WT and TKO signals were multiplied by a scale factor of 2, to facilitate visualization in a single plot. ChIP-seq data are from Geeven et al. (2015). **(d)** Go-term enrichment analysis of the set of genes proximal to a differential lncRNA. −log(p-value) for each term is plotted. **(e)** IGV browser snapshots of lncRNAs derived from the annotated enhancers regulating *Nanog* (upper panel) and *Myc* (lower panel). **(f)** Heatmap showing the expression fold-changes of upregulated or downregulated lncRNAs and the neighbouring coding gene.

### Depleting histone H1 levels in human cells triggers non-coding transcript accumulation in chromatin and transcription-dependent replicative stress

To confirm that the accumulation of non-coding transcripts in chromatin was a direct consequence of histone H1 depletion, rather than an indirect effect related to the lack of differentiation potential of H1-TKO mESCs (28), we next analyzed the transcriptional status of human differentiated cells knocked-down for histone H1 (19). We applied the same computational pipeline designed for CheRNA-seq data to re-analyze published total RNA-seq data from breast cancer T47D cells upon doxycycline-induced knock-down for the subtypes H1.2 and H1.4 (shMultiH1, human homologues of the murine *H1c* and *H1e*, respectively) (**Supplementary Figure 2a**; 19). Despite the reduced representation of noncoding transcripts in total RNA preparations relative to chromatin-RNA preparations, lncRNA transcription was enhanced upon induction of histone H1 silencing (**Figure 3a**). These lncRNAs were enriched in CheRNA preparations relative to nucleoplasm (**Supplementary Figure 2b**), and the genomic loci around their TSS were marked by H3H4me1 (**Supplementary Figure 2c**, left panel). In addition, both the levels of H3H4me3 and the chromatin accumulation of differential lncRNAs increased upon histone H1 depletion (**Supplementary Figure 2b-c**).

**Figure 3.**
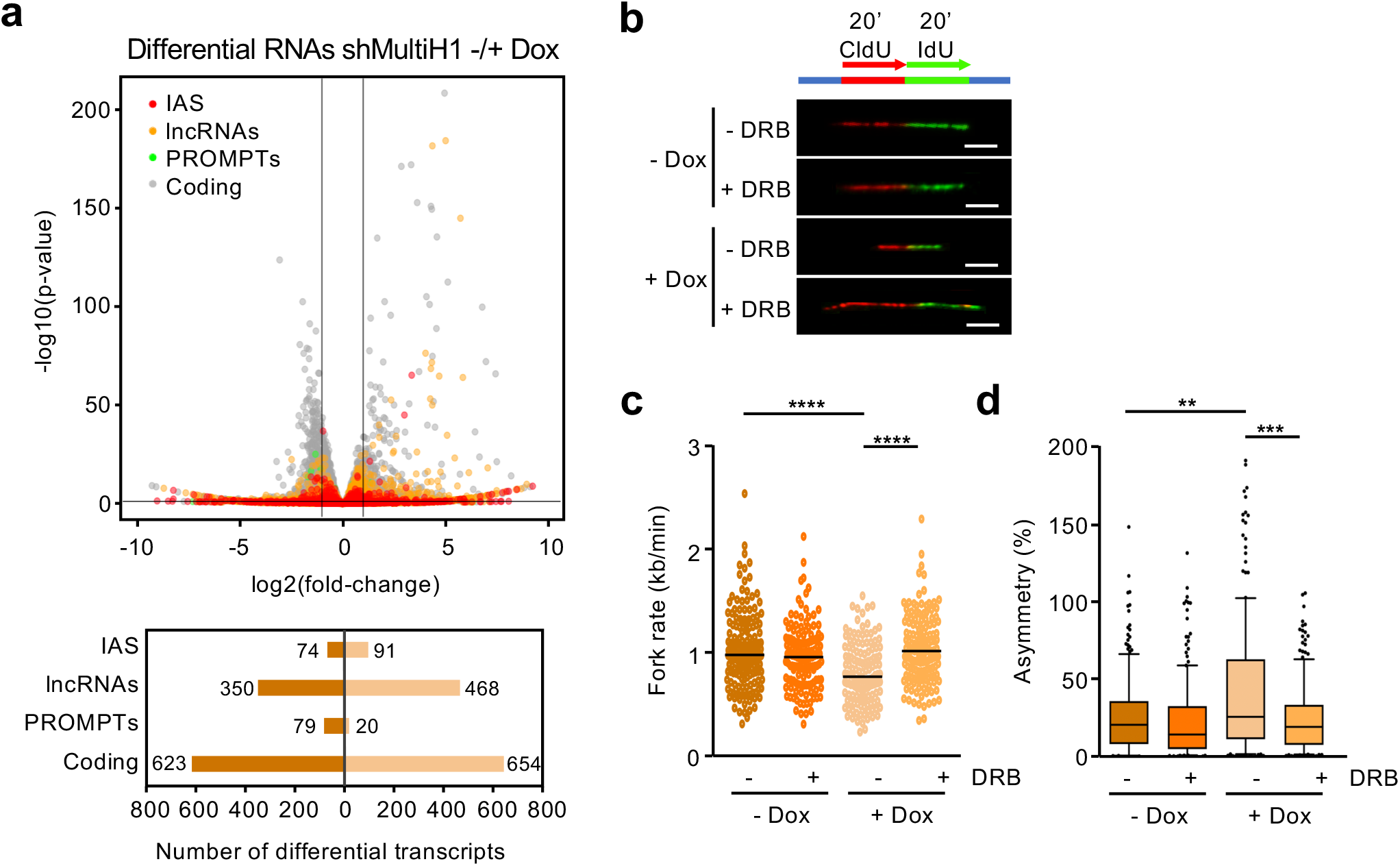
lncRNAs overexpression and transcription-dependent replicative stress upon inducible knock-down of histone H1 in human cells. **(a)** Volcano plot showing the log2(fold-change) and the −log(p-value) for each transcript in shMultiH1 TD47 Dox+ cells relative to Dox-cells. The number of differential expressed transcripts for each category is shown below. Total RNA-seq data are from Izquierdo-Boulstridge et al (2017). **(b)** Representative example of DNA fibers from Dox- and Dox+ shMultiH1 cells labelled sequentially for 20 min with CldU (red) and IdU (green) in conditions of active (-DRB) or blocked (+DRB) transcription elongation used to estimate fork rates and fork asymmetries. Scale bar, 5 μm. Measure of DNA replication fork rate **(c)**, and fork asymmetry **(d)**, in Dox- and Dox+ shMultiH1 cells untreated (-DRB) or treated for 3h with DRB (+DRB). Median values are indicated. Data shown are pooled from two independent experiments, n > 135. Statistical differences between distributions were assessed with the Mann-Whitney rank sum test. p-value: **<0.01; ***<0.001; ****<0.0001.

We then asked whether this short-term histone H1 depletion also recapitulate the replicative stress we described for H1-TKO mESCs (14). DNA fiber analysis showed significant decreases in fork rate and increases in fork asymmetry upon H1 reduction (**Figure 3b-d**). Most importantly, the replicative phenotypes were transcription dependent, as both were readily reverted when inhibiting RNAPII elongation activity by 5,6-dichlorobenzimidazole1-β-D-ribofuranoside (DRB) treatment. The elevated DNA damage signalling of these cells was concomitantly reduced by transcription inhibition (19 and **Supplementary Figure 2d**). All together, these results indicate that, as in H1-TKO mESCs, reducing histone H1 content in human differentiated cells increases non-coding RNA chromatin association and transcription-dependent replicative stress.

### Accumulated transcripts are tethered to chromatin by RNAPII and are transcribed at high rates

To address how non-coding RNAs were transcribed we next investigated RNAPII genomic occupancies. We performed chromatin immunoprecipitation sequencing (ChIP-seq) using human chromatin as a spike-in control to detect quantitative differences in chromatin-bound RNAPII between cell types (see Methods), and found a 12% more RNAPII in H1-TKO chromatin (**Supplementary Figure 3a**). However limited, this excess in RNAPII complexes was not uniformly distributed through the genome, but specifically located around the TSS of accumulated transcripts in H1-TKO chromatin (**Supplementary Figure 3b-c**). Changes in the levels of coding transcripts, lncRNAs and IAS were accompanied by parallel changes in the levels of RNAPII at their TSS in either cell type (**Figure 4a**). Moreover, promoter-proximal regions of lncRNAs and IAS upregulated in H1-TKO cells recruit almost as much RNAPII as some coding promoters (**Figure 4a**-lower panels). To ensure that differential transcripts anchored to chromatin by RNAPII complexes were actively transcribed, we conducted transient transcriptome sequencing assays (TT-seq; 29). We found a positive correlation between RNA synthesis rates and chromatin transcript abundances for all RNA classes, indicating that the residence time of nascent transcripts on chromatin is related to their production (**Figure 4b-c**). Collectively, these data suggest that histone H1 prevents the recruitment of RNAPII complexes driving transcription of non-coding RNAs.

**Figure 4.**
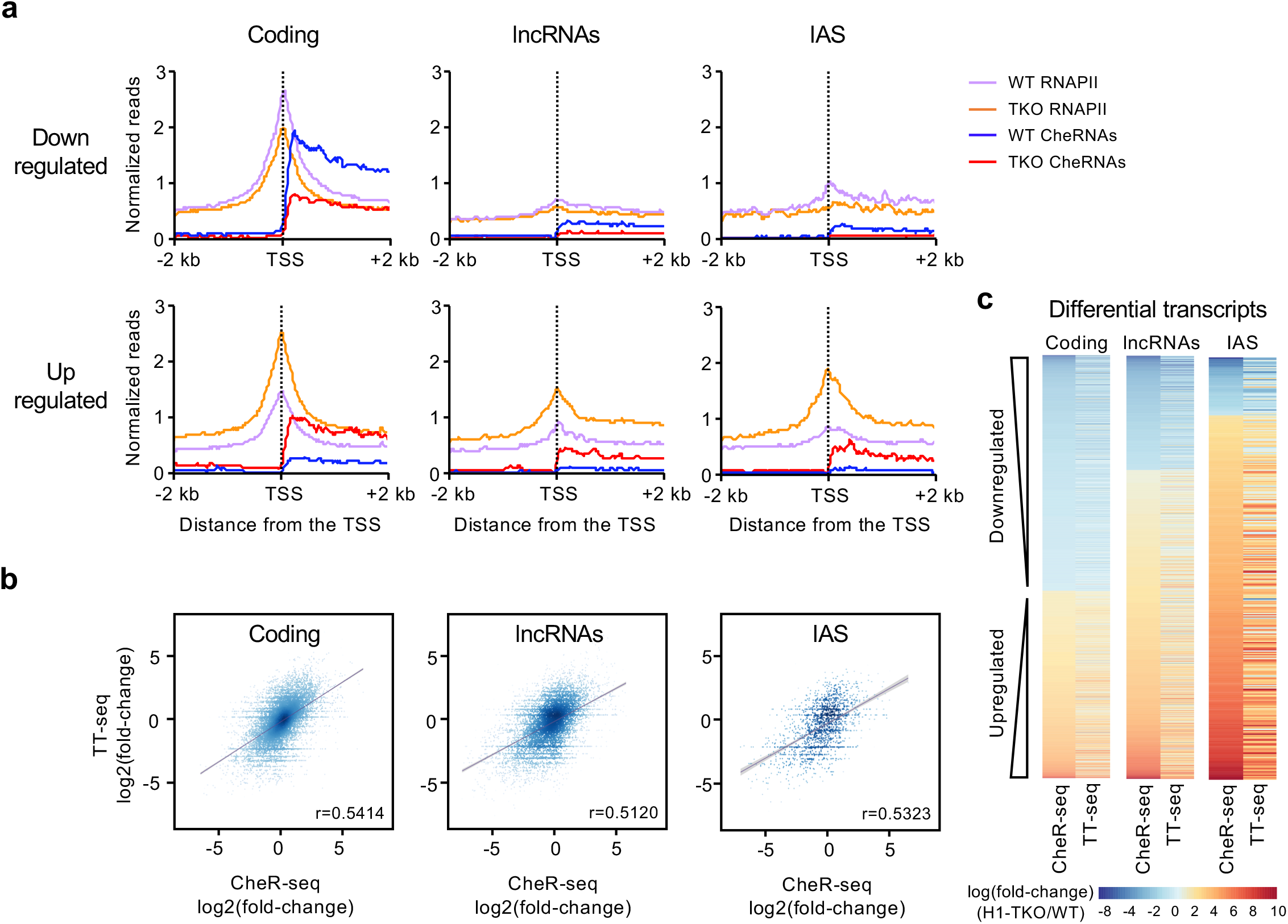
Accumulated transcripts are transcribed at high rates and tethered to chromatin by RNAPII. **(a)** Metaplots of RNAPII-seq and CheRNA-seq spike-in normalized signals at the indicated transcript categories in WT and H1-TKO cells, plotted in a 4kb window around the TSS. RNAPII WT and TKO signals were multiplied by a scale factor of 1:2, to facilitate the visualization in a single plot. **(b)** Relationship between expression fold-changes (CheR-seq) and transcriptional rate fold changes (TT-seq) at the indicated transcript categories. **(c)** Heatmap representation of CheRNA-seq and TT-seq fold-changes between H1-TKO/WT conditions at differential transcripts.

### Chromatin-associated transcripts have reduced levels of m6A modification

Chromatin-associated regulatory RNAs, including PROMPTs and enhancer RNAs, are marked by N6-methyladenosine (m6A) modification, what destabilizes their levels in chromatin (30). Thus, in order to identify specific features of lncRNA transcripts repressed by H1, we first focused on m6A. This modification is co-transcriptionally deposited on RNAPII transcripts by the METTL3/METTL14 writer complex and read by the YTH-domain-containing proteins (31). Intriguingly, several members of the writer complex accessory proteins, as well as the m6A nuclear reader YTHDC1, have been identified as high-confidence interactors of multiple histone H1 subtypes in human cells by proteomics approaches (32). We hypothesized that non-coding transcripts in H1-TKO cells might accumulate in chromatin due not only to an increased transcription (**Figure 4**), but also to alterations in m6A modification. In agreement with this idea, global m6A levels were decreased on CheRNA preparations but not on total RNA in H1-TKO cells relative to their WT counterparts (**Figure 5a**). To determine m6A changes at specific transcript classes we coupled CheRNA purification with m6A immunoprecipitation and sequencing (ChMeRIP-seq), using an *in vitro* methylated RNA as spike-in control (see Methods). ChMeRIP reads distribution across gene bodies recapitulated the TSS- and stop codon-proximal accumulation reported by MeRIP-seq from total RNA in mESCs (30, 33, 34) (**Supplementary Figure 4a-b**). It also detects m6A enrichments at characterized lncRNAs, as exemplified by *Neat1* or *Malat1* loci (33, 35) (**Supplementary Figure 4c**). As anticipated from the global m6A CheRNA measurements, ChMeRIP-seq analyses revealed significantly reduced levels of methylation in H1-TKO cells for both coding and non-coding transcripts (**Figure 5b** and **Supplementary Figure 4a** and **4d**). To note, lncRNAs displayed higher m6A levels than coding RNAs in WT mESCs chromatin (**Figure 5b**), a finding suggestive of a distinct post-transcriptional regulation dynamics for non-coding transcripts. Interestingly, the H1-TKO/WT fold-change in m6A levels was higher for lncRNAs, especially for the upregulated lncRNA category (**Figure 5c**, and representative examples in **Figure 5d**), which consistently harbor higher binding densities of the m6A catalytic subunit METTL3 at their promoter regions in WT mESCs (**Supplementary Figure 4e**; data from 34). To further characterize the transcripts mostly affected by m6A loss, we sorted them by nascent RNA abundances and histone H1 promoter-proximal occupancies. We found that maximal reductions in m6A levels correlated with lower transcript expression and higher histone H1 occupancy in WT cells (**Figure 5e-f**). As noted before, these associations were stronger for non-coding transcripts (**Figure 5e-f**, lower panels), albeit also occur at coding RNAs (**Figure 5e-f**, upper panels). Altogether, these results provide evidence supporting a connection between histone H1 and m6A levels of chromatin-associated lncRNAs.

**Figure 5.**
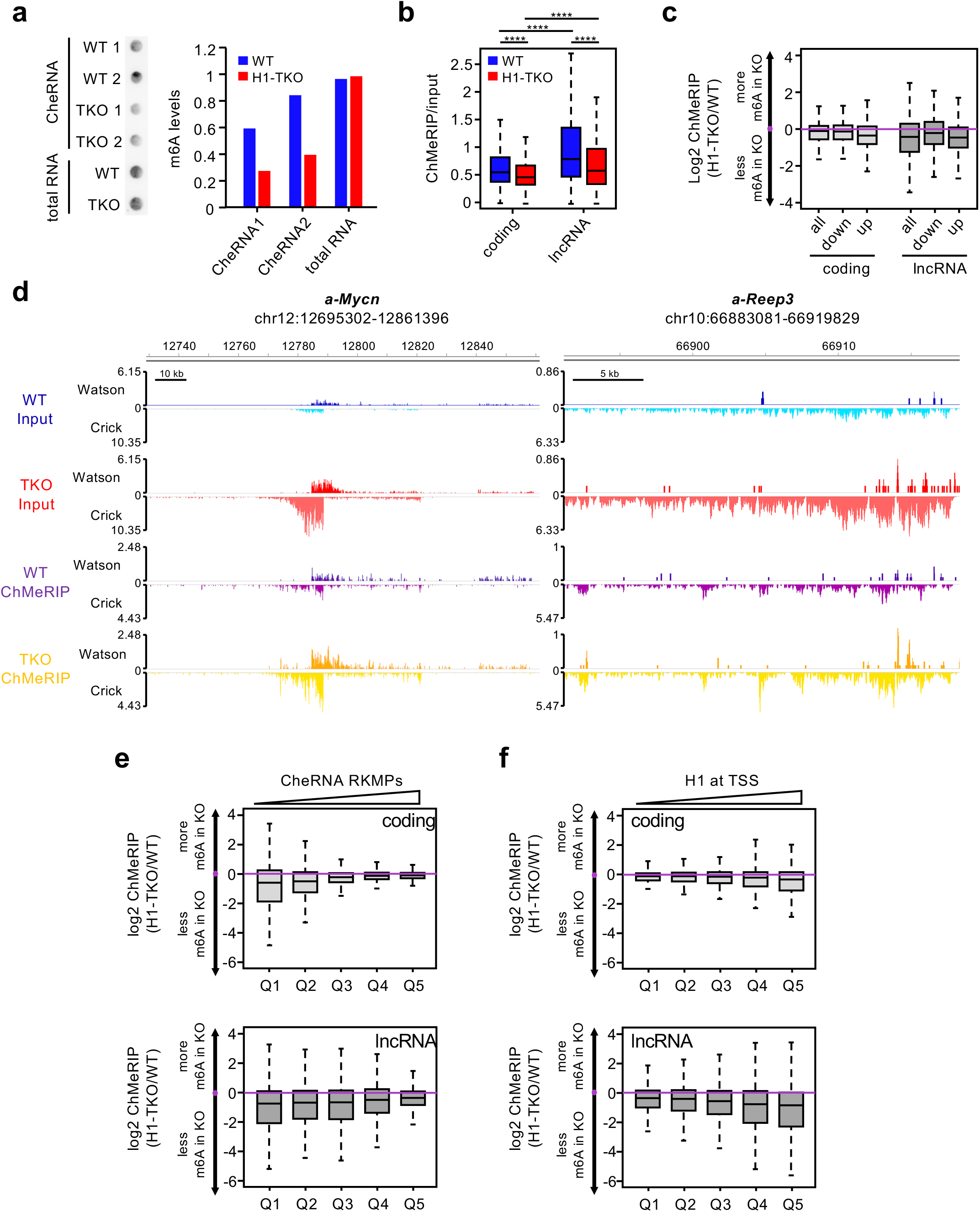
Accumulated lncRNAs in H1-TKO cells have reduced levels of m6A modification. **(a)** Dot-blot quantification of m6A levels on 5 □g of CheRNA or total RNA preparations. (**b**) ChMeRIP RPKMs for coding and non-coding transcripts, normalized by the RPKMs in the CheRNA input. ****Mann-Whitney-Wilcoxon test p-value<2.2×10^-16^ for all comparisons. (**c**) Plots showing the ratios of normalized ChMeRIP read counts relative to input between H1-TKO and WT cells at the indicated transcript categories. **(d)** Representative IGV screenshots showing normalized reads from ChMeRIP-seq and the corresponding CheRNA-seq inputs of two differentially abundant lncRNAs in WT and H1-TKO cells. (**e**) ChMeRIP ratios between H1-TKO and WT cells across 5 quantiles of increasing chromatin transcript levels for coding (upper plot) or lncRNA (lower plot). (**f**) Same as in (e) across 5 quantiles of increasing histone H1 levels at -/+ 2 kb of the TSS. H1 ChIP-seq data are from Cao et al (2013). In all cases boxplots denote the medians and the interquantile ranges and the whiskers represent the 10 and 90 percentiles.

To sustain these findings further, we next examined m6A and chromatin RNA abundances of H1-TKO differentially expressed RNAs in mESCs depleted of METTL3 (*Mettl3-KO*) (30). As expected, m6A levels were reduced in *Mettl3*-KO cells, irrespectively of the transcript category analyzed (**Supplementary Figure 4f**, empty bars). Notably, we identified a positive correlation between expression alterations in both mutant scenarios: up or downregulated lncRNAs in H1-TKO cells were also up or down regulated in *Mettl3-KO* cells (**Supplementary Figure 4g**, empty bars). These observations suggest that, while affecting similarly all types of transcripts, reduced m6A levels specifically stabilize non-coding transcripts that are retained in chromatin in histone H1-depleted cells. Similar analyses in cells depleted of the m6A nuclear reader YTHDC1 revealed that RNA abundances were negatively correlated in all cases between histone H1- or METTL3-depleted and YTHDC1-depleted cells (**Supplementary Figure 4g**, grey bars), suggesting that YTHDC1 is implicated in the differential stability of these non-coding transcripts. Supporting a link between components of the m6A pathway and histone H1, all somatic subtypes (H1a, H1b, H1c, H1d, H1e), and the testicular isoform H1t, were strongly up-regulated in *Mettl3-KO* cells (**Supplementary Figure 4h**).

### Reduced methylation of lncRNAs alters their chromatin turnover triggering replication-transcription conflicts

Taken together, these analyses indicate that reductions in histone H1 levels results in the altered turnover of non-coding transcripts in chromatin in a m6A-dependent manner. This interpretation makes the prediction that increasing m6A levels in H1-TKO cells will leverage the retention of non-coding transcripts in chromatin, thus alleviating the replicative stress phenotype. To test this prediction we used reversible short interfering RNA (siRNA) to deplete the m6A erasers, ALKBH5 and FTO (**Figure 6a**, left and central panels). Then, we analyzed replication dynamics by fiber stretching. Interestingly, the depletion of either demethylase restored the replication fork speed of H1-TKO cells towards WT levels (**Figure 6b**). Analogous effects were observed by a combined depletion of both demethylases (**Supplementary Figure 5a-b**), or by exposing the cells to the alpha-ketoglutarate dependent demethylases inhibitor R-2-hydroxyglutarate (R2-HG) in conditions in which overall DNA synthesis was not detectably altered (**Supplementary Figure 5c-d**). Intriguingly, the replicative phenotype was also fully restored upon knockdown of the m6A reader YTHDC1 (**Figure 6b**, rightmost panel), even though its expression was only decreased by 30% in H1-TKO cells (**Figure 6a**). These results imply reduced methylation of non-coding RNAs as a trigger of replication-transcription conflicts in H1-depleted cells.

**Figure 6.**
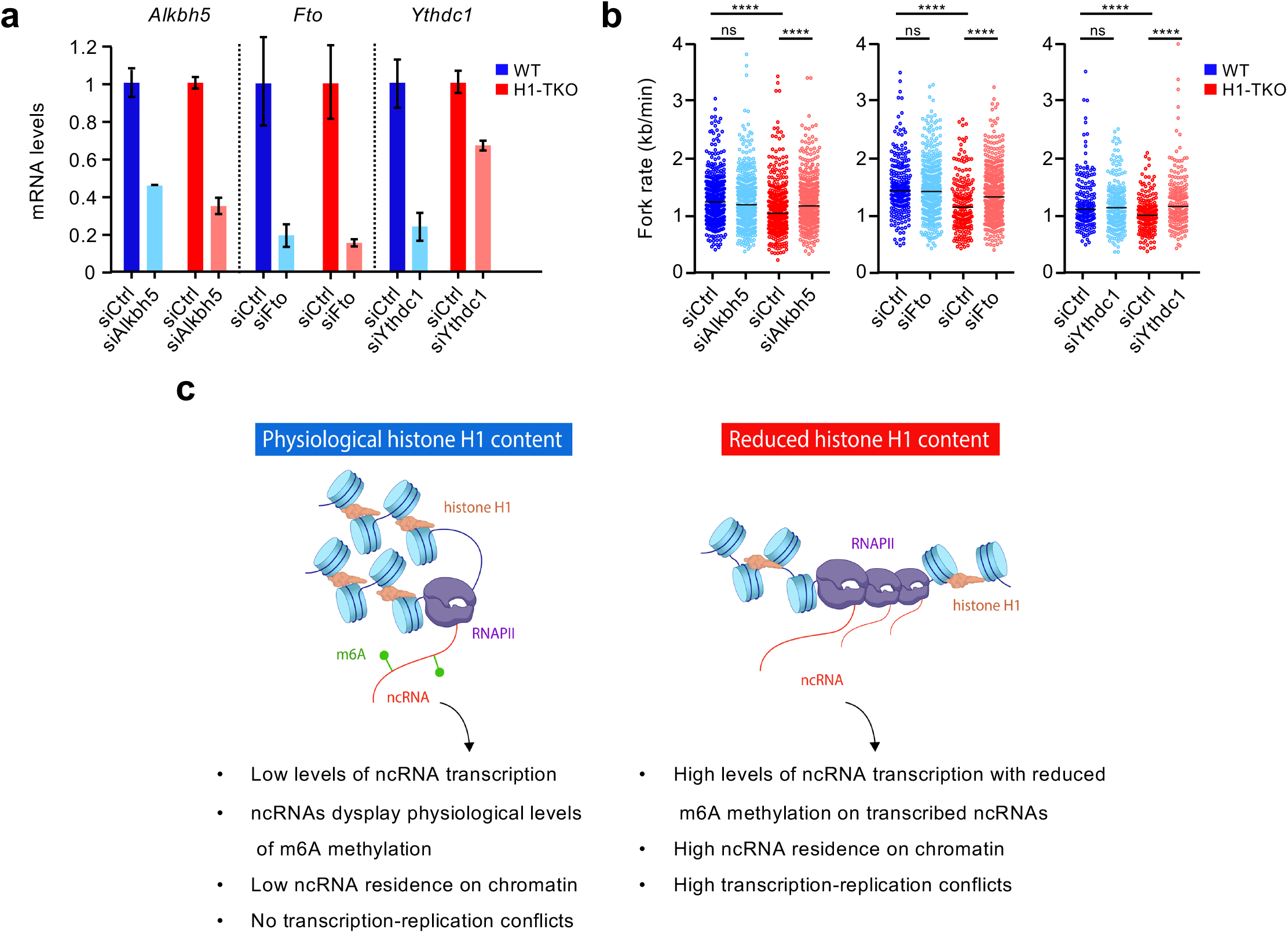
Impairing m6A demethylase activity in H1-TKO cells decreases lncRNA abundance on chromatin and rescues the speed of replication forks. **(a)** RT-qPCR mRNA silencing levels of *Alkbh5*, *Fto* and *Ythdc1* 24h after cellular transfection with specific siRNAs. mRNA levels were normalized to *Hprt* mRNA levels at each condition. Data show the median +/-s.d of two independent replicates (n=2). **(b)** Replication fork rates of WT and H1-TKO cells transfected with the indicated siRNAs. Median values are indicated. Data shown are pooled from two independent experiments, n > 410. Statistical differences between distributions were assessed with the Mann-Whitney rank sum test. p-value: ****<0.0001. (**c**) Cartoon representing the impact of histone H1 levels on ncRNA turnover on chromatin. Under physiological H1 levels, transcription of ncRNAs is reduced and those present display high levels of m6A methylation. Upon H1 depletion, there is an increased recruitment of RNAPII complexes driving transcription of ncRNAs. These nascent ncRNAs, in addition, have reduced levels of m6A modification, what causes their stabilization on chromatin generating conflicts with incoming DNA replication forks.

## DISCUSSION

In all organisms studied, reductions in H1 levels do not cause global upregulation of transcription but rather affect a reduced set of genes (5, 7, 15–18, 36). By examining nascent transcription and RNAPII occupancies in WT and H1-TKO mES cells we expanded these findings to around 1,300 upregulated and 1,600 downregulated genes, mostly implicated in cell differentiation and development (**Supplementary Figure 6a-d**). Altered H1 content likely affects RNAPII recruitment at those TSS sites directly, as the epigenetic architecture of the promoter-proximal region of differentially expressed genes was not significantly changed (**Supplementary Figure 6e**). We further show that H1 depletion allowed increased recruitment of RNAPII complexes driving transcription of thousands of non-coding RNAs, 75% of which were not previously annotated in the mouse genome (ENSEMBL non-coding transcriptome). Strikingly, we found that lncRNAs harbor reduced levels of m6A modification, causing their stabilization on chromatin and thus generating conflicts with advancing replication forks (**Figure 6c**). These findings are highly relevant as they uncover a novel link between an essential component of chromatin and the m6A non-coding regulatory axis, with important implications for genome integrity.

Although m6A modification is best-studied in coding RNAs, m6A profiling studies have shown that it is also present in lncRNAs (37). Methylation of *XIST* lncRNA contributes to its transcriptional repression effects through the recruitment of the nuclear m6A binding protein YTDC1 (35), although the extent of its contribution to *XIST*-mediated chromosomal silencing remains controversial (38). The question of how m6A modification is selectively directed to specific RNAs is not yet clear. It has been proposed that the RNA binding proteins RBM15 and RBM15B, that associate with the WTAP-METTL3/14 complex, enable the binding of the m6A writer complex to multiple RNAs, including *XIST* (35). In a similar fashion, the components of the WTAP complex VIRMA (virilizer homolog), WTAP (Wilms Tumor Associated Protein), ZC3H13, and CBLL1/Hakai, all of which interact with histone H1 (28), are putative candidates to mediate substrate RNA specificity. Since reductions in m6A levels correlate positively with histone H1 occupancies in WT cells (**Figure 5f**), we speculate that H1 local abundances at lncRNAs loci will not only limit RNAPII recruitment at their TSS sites, but also facilitate co-transcriptional m6A installation thus ensuring appropriate non-coding transcript turnover. m6A-RNA turnover dynamics are executed by reader proteins, and several mechanisms have been proposed depending on the RNA type and cellular context (39, 40). Conditional knockout of YTHDC1 in mESCs, for example, enhance the stability of repeat RNAs transcribed by transposons (30), and initiates cellular reprogramming to a 2C-like state (33). On the other hand, YTHDC1-m6A RNAs can form phase-separated nuclear condensates that maintain mRNA stability suppressing myeloid leukemic differentiation (41). The intriguing finding that YTHDC1 depletion in H1-TKO cells rescues the replicative phenotype similarly to ALKBH5/FTO depletion suggests that YTHDC1 is required for the stability of lncRNAs even when m6A levels are impaired. This is in agreement with a recent preprint showing that YTHDC1 mediates the chromatin association and gene expression effects of HOTAIR lncRNA regardless the ablation of its major m6A site (42). The authors propose that differential affinities of YTHDC1 for distinct m6A sites might mediate the functionally diverse and context dependent effects observed even for a single lncRNA. Further experiments are required to determine both the mechanisms of H1-mediated m6A deposition and m6A-mediated fate at individual lncRNAs.

Our work highlights yet another example of the multi-faceted functions of histone H1 beyond chromatin architecture. We propose that histone H1 functions as a regulator of lncRNA metabolism. These findings add onto the hypothesis that the multi-functionality of linker histones can be explained through H1-protein interactions that directly regulate recruitment of proteins to chromatin (1, 43). The adaptability of the intrinsically disordered N- and C-terminal domains of H1 likely facilitates the wide range of specific protein-protein interactions reported (32, 44, 45). Accordingly, the disordered terminal domains acquire secondary structure when bound to DNA (46), nucleosomes (47, 48), and possibly other proteins. Recent studies addressing histone H1 distribution in the three-dimensional nucleus showed that, in differentiated cells, local H1 density regulates the degree of chromatin compaction through maintaining a condensed and spatially distinct chromatin B compartment (49, 50). In line with this, decompaction of three-dimensional chromatin has been proposed as the dominant effect of H1 loss of function occurring in B cell lymphomas (51). Given the implication of histone H1 in lncRNA modification and chromatin retention unveiled here, it is tempting to speculate that some of the defects associated with diseases carrying histone H1 missense mutations, such as certain cancers or Rahman syndrome (52–54), could be mediated by lncRNAs altering the maintenance of proper chromatin compartmentalization, or nuclear bodies formation (55, 56).

In conclusion, our work emphasizes the crucial role of histone H1 as an epigenetic controller, whose full characterization awaits multiple studies in the coming years. As anticipated by A. P. Wolffe, “understanding the molecular mechanisms by which histone H1 exerts its functions might uncover new ways to manipulate gene expression” (57).

## METHODS

### Cell culture, siRNA transfection and drug treatments

Mouse embryonic stem cells were grown in DMEM (Invitrogen) supplemented with 10% fetal bovine serum (Biosera), 1x non-essential aminoacids (Gibco), 1mM sodium piruvate (Gibco), 2mM L-glutamine (Gibco), 50 μM β-mercaptoethanol (Gibco), 10^3^ U/mL LIF (ESGRO), 100 U/mL penicillin (Invitrogen) and 100 μg/mL streptomycin (Invitrogen), at 37°C and 5% CO_2_. For transcription inhibition experiments, cells at 80% confluency were treated with 100 μM 5,6-dichlorobenzimidazole 1-b-d-ribofuranoside (DRB) (Sigma) for 3 hours. For small interfering RNAs (siRNA) transfection, Lipofectamine™ 2000 (Thermo) was used to deliver siRNAs into mES cells. 40nM of each siRNA was diluted in 1ml of OPTIMEM and incubated for at least 20 minutes with Lipofectamine diluted in OPTIMEM. Cells were tripsinized, resuspended in medium without antibiotics and added to the previous mix of Lipofectamine and siRNA. After incubating cells for 15 min at RT, cells were seeded into new plates. Transfected cells were analyzed after 24h. All siRNAs were purchased from Sigma and their sequences are listed in **Supplementary Table 1**. For FTO/ALKBH5 inhibition, cells were treated with 20 or 40 mM R-2-hydroxyglutarate (R2-HG) (Sigma) for 24h. T47D inducible H1 knock-down cell lines were grown in RPMI 1640 medium, supplemented with 10% FBS, 2 mM L-glutamine, 100 U/ml penicillin, and 100 mg/ml streptomycin at 37°C with 5% CO2. Depletions were induced by 3 days exposure to 2.5 μg/ml doxycycline (Sigma) as described (19). All cells tested negative for mycoplasm infection.

### Flow Cytometry

For cell-cycle analyses, cells were pulse-labeled for 20 minutes with 250 μM IdU (Sigma) and fixed overnight in 70% ethanol at −20°C. Cells were then incubated in 2 M HCl (Merck) with 0.5% Triton X-100 (Sigma) for 30 minutes and neutralized with 0,1 M Sodium Tetraborate pH 9.5 (Merck) for 2 minutes before blocking 10 minutes with a solution of 1% BSA (Sigma) and 0.5% Tween20 (Sigma) in PBS. Cells were then incubated for 1 hour with mouse anti-BrdU antibodies (BD Biosciences) followed by 30 minutes incubation with anti-mouse IgG Alexa-Fluor 647 antibodies (Thermo Fisher Scientific) at RT. Cells were finally stained either with 2μg/ml DAPI (Merck) or with PI/RNAse cycle buffer (BD Pharmingen) for another 30 minutes in the dark at RT. All samples were processed in a FACSCanto II (Becton Dickinson) with FACSDiva v6.1.3 analysis software and analysed with the FlowJo v10 program.

### Chromatin enriched RNA (CheRNA) sequencing

CheRNA preparations were obtained as described in (20). Briefly, 40×10^6^ mES cells were lysed in 800 μL ice-cold Lysis Buffer A (10 mM Tris pH 7.5, 0.1% NP-40, 150 mM NaCl) for 5 minutes on ice. Nuclei were collected by sucrose cushion centrifugation (24% sucrose in lysis buffer A), rinsed with ice-cold PBS + 1mM EDTA, and resuspended in 500μL ice-cold Glycerol Buffer (20 mM Tris pH 7.9, 75 mM NaCl, 0.5 mM EDTA, 0.85 mM DTT, 0.125 mM PMSF, 50% glycerol). Nuclei were lysed by adding one volume of ice-cold Lysis Buffer B (10 mM HEPES pH 7.6, 1mM DTT, 7.5 mM MgCl2, 0.2 mM EDTA, 0.3M NaCl, 1M urea, 1% NP-40) and kept on ice for 10 minutes, with periodic vigorous shaking. Insoluble chromatin was sedimented by centrifugation at 14000g and 4°C for 2 minutes, rinsed twice with cold PBS + 1mM EDTA, and resuspended in 100 μL PBS. At this point, 10 pg of an in vitro transcribed luciferase RNA was added to both the nucleoplasmic and chromatin samples as a spike-in control. RNAs were purified using TRIzol™, following manufacturer’s instructions. Before library preparation, ribosomal RNA was depleted from the samples by a treatment with Ribo-Zero rRNA Removal Kit (Illumina). Libraries were generated using TruSeq Stranded Total RNA Library Prep (Illumina), and sequenced by 1×75 single reads at the Fundación Parque Científico de Madrid.

### DNA Fiber stretching

Exponentially growing cells were pulsed consecutively for 20 minutes with 50 mM CldU (Sigma) and 250 mM IdU (Sigma). Collected cells were resuspended in cold PBS at a concentration of 0.5×10^6^ cells/mL, and 2μL of this cell suspension was lysed through the addition of 10 μL of spreading buffer (200 mM Tris pH 7.4, 50 mM EDTA, 0.5% SDS) on the top of a microscopy slide at 30°C. After 6 minutes of incubation in a humidity chamber at RT, DNA fibers were stretched by leaning the slide with a 30° slope. Slides were air dried and fixed at −20°C with 3:1 methanol:acetic acid solution, incubated with 2.5M HCl solution for 30 minutes at RT, washed three times with PBS, and treated with blocking solution (1% BSA, 1% Triton X-100 in PBS) for 1 hour. Samples were sequentially incubated with primary antibodies; 1:100 anti-CldU (Abcam), 1:100 anti-IdU (Bencton Dickinson) and 1:3000 anti-ssDNA (Millipore) for one hour, and with secondary antibodies; 1:300 anti-rat IgG Alexa-Fluor 594, anti-mouse IgG1 Alexa-Fluor 488 and antimouse IgG2a Alexa-Fluor 647 (Invitrogen) for 30 minutes. Slides were mounted with Prolong Diamond (Invitrogen) and fibers visual acquisition was performed with an Axiovert200 Fluorescence Resonance Energy Transfer microscope (Zeiss) using the 40x oil objective. Images were analyzed with ImageJ software, considering a conversion factor of 1μm=2.59 kb (58). Fork rates were calculated by measuring the length (kb) of the IdU track divided by the duration of the pulse (min), and fork asymmetries were obtained by calculating the percentage of the difference between the lengths of both CldU and IdU tracks of each replication fork. Statistical analysis of all data was performed with Prism v5.0.4 (GraphPad Software) using the non-parametric Mann-Whitney rank sum test.

### Immunofluorescence

Cells grown on glass coverslips (VWR) were fixed with 3.7% formaldehyde in PBS for 15 minutes at RT and permeabilized with 0.5% Triton X-100 in PBS for 20 minutes at RT. Samples were blocked with 3% BSA (Sigma) in PBS before overnight incubation at 4°C with antibodies anti-γH2AX (1:250) (Millipore), followed by 1 hour incubation at RT with antibodies anti-rabbit Alexa-Fluor 488 (Invitrogen) and 5 minutes staining at RT with 2ng/μl of DAPI (Merck) in PBS. Coverslips were mounted in Prolong Diamond (Life Technologies) and visual acquisition was performed in a A1R+ confocal microscope (Nikon) using a either a 40x or a 60x oil objective. Nuclear segmentation was based on DAPI staining. Statistical analyses were performed in Prism v5.0.4 (GraphPad Software) using the non-parametric Mann-Whitney rank sum test.

### Chromatin immunoprecipitation and quantitative real time PCR (ChIP-qPCR)

Chromatin immunoprecipitation was performed according to the Upstate (Millipore) standard protocol. Briefly, cells were fixed using 1% formaldehyde for 10 min at 37°C, chromatin was extracted and sonicated to generate fragments between 200 and 500 bp. Next, 30 μg of sheared chromatin was immunoprecipitated overnight with anti-H3K4me3 (Abcam), anti-H3K4me1 (Abcam) or Rabbit IgG (Millipore) antibodies. Immunocomplexes were recovered using 20 μl of protein A magnetic beads, washed and eluted. Cross-linking was reversed at 65°C overnight and immunoprecipitated DNA was recovered using the IPure Kit (Diagenode). Genomic regions of interest were identified by realtime PCR (qPCR) using SYBR Green Master Mix (Invitrogen) on a QuantStudio™ 5 Real-Time PCR System machine (ThermoFisher Scientific). Each value was normalized by the corresponding input chromatin sample. Oligonucleotide sequences are detailed in **Supplementary Table 1**.

### Retrotranscription and quantitative real time PCR (RT-qPCR)

Superscript III (Invitrogen) was used to generate the cDNA following manufacturer’s instructions. qPCR reactions were performed in an ABI Prism 7900HT Detection System (Applied Biosystems), using HotStarTaq DNA polymerase (Qiagen) following manufacturer’s instructions. For absolute quantification, the Ct of each amplicon was interpolated in a standard curve obtained from the amplification of genomic DNA at five different concentrations (from 0.2ng/μL to 125ng/μL) and analyses were carried out with the SDS2.4 software (Applied Biosystems). Primer sequences are listed in **Supplementary Table 1**.

### RNAPII chromatin immunoprecipitation sequencing (ChIP-seq)

Crosslinkings were performed in culture medium with 1% formaldehyde during 15 minutes at RT. After stopping the reactions with 125 mM glycine for 5 minutes, cells were washed twice with PBS and collected by scrapping in ice-cold PBS supplemented with protease and phosphatase inhibitors (10 μM leupeptin, 100 μM PMSF, 1μM pepstatin, 2 μg/mL aprotinin, 5 mM NaF, 1mM NaVO3). Cells were centrifuged at 200g for 5 minutes, resuspended in cold Lysis Buffer (50 mM Tris pH 8, 1% SDS, 10 mM EDTA, protease and phosphatase inhibitors) at a concentration of 2×10^7^ cells/mL and incubated on ice for 20 minutes. Soluble chromatin was fragmented on a Covaris sonication system by 40 cycles at 20% intensity, during 20 minutes. 100 μg of the fragmented chromatin was diluted 1:10 in Dilution Buffer (20 mM Tris pH 8, 1% Triton X-100, 2mM EDTA, 150mM NaCl, protease and phosphatase inhibitors), and 5 μg of human chromatin, obtained from a MCF10A cell line following the same protocol, was added as spike-in control. Precleared chromatin was incubated overnight with 25 μg of anti-RNApolII antibody (Millipore) at 4°C with gentle agitation, followed by a 2 hours incubation with 200 μL of A/G protein beads. Immunocomplexes were washed sequentially with four different buffers, all supplemented with protease and phosphatase inhibitors: low salt buffer (10 mM Tris pH 8, 0.1% SDS, 1% Triton X-100, 2mM EDTA, 150 mM NaCl), high salt buffer (20 mM Tris pH 8, 0.1% SDS, 1% Triton X-100, 2mM EDTA, 500 mM NaCl), LiCl buffer (10 mM Tris pH 8, 0.25M LiCl, 1% NP40, 1% Na-deoxycholate, 1mM EDTA) and TE buffer (10 mM Tris pH 8, 1mM EDTA). Finally, chromatin was eluted with elution buffer (0.1M NaHCO3, 1% SDS), crosslinkings were reverted, and DNA was purified with phenolchloroform extraction and ethanol precipitation. Libraries were generated with the NEBNext Ultra II DNA Library Prep Kit (New England Biolabs) following the manufacturer’s recommendations and sequenced by 1×75 single-reads at Fundación Parque Científico de Madrid.

### Transient transcription sequencing (TTseq)

Nascent transcription labeling assays were carried out as previously described (29, 59). Briefly, 4-thiouridine (4sU) was added to sub-confluent cell cultures at a final concentration of 1 mM for 10 min before cell harvest. Cells were lysed directly on a plate with 5 ml of TRIzol (Invitrogen), total RNA was isolated following manufacturer’s protocol and sonicated by two pulses of 30 seconds in a Bioruptor instrument. A total of 100 μg of sonicated RNA per cell line was used for biotinylation and purification of 4sU-labeled nascent RNAs. Biotinylation reactions consisted of total RNA and EZ-Link HPDP-Biotin dissolved in dimethylformamide (DMF) and were performed in labeling buffer (10 mM Tris pH 7.4, 1 mM EDTA) for 2 h with rotation at RT. Unbound Biotin-HPDP was removed by chloroform/isoamylalcohol (24:1) extraction in MaXtract tubes (Qiagen). RNA was precipitated with 10^th^ volume of 5M NaCl and 1 volume of isopropanol. Following one wash in 80% ethanol, the RNA pellet was left to dry and resuspended in 100 μL RNase-free water. Biotinylated RNA was purified using □ Macs Streptavidin kit, eluted twice using 100 mM DTT and recovered using RNeasy MinElute Cleanup column (Qiagen) according to instructions. cDNA libraries were prepared using NEBNext Ultra Directional RNA Library Prep Kit (New England Biolabs) according to the manufacturer’s instructions. Libraries were pooled and sequenced by 2×75 single-reads at Fundación Parque Científico de Madrid. Reads were aligned to the mm10 reference genome using Tophat2 (60) with standard parameters. bedGraph files loaded in the IGV browser were generated with the Bedtool genomecov.

### CheRNA m6A-RNA immunoprecipitation sequencing (ChMeRIP-seq)

CheRNAs were purified as described above, and 1 pg of *in vitro* transcribed N6-methylated Luciferase RNA per million cells was added as spike-in control before Trizol extraction. RNA fragmentation and meRIP were performed as described (61, 62), with the following modifications. Aliquots containing 2 μg of CheRNAs in 18μL of DEPC H2O were fragmented by incubation with 2 μL of 100 uL Tris pH 7, 100 uL ZnCl2 1M, 800 uL DEPC water, at 70°C for 8 minutes, and reactions were stopped by adding 2 μL of 0.5M EDTA. A total of 50 μg of pooled fragmented CheRNAs were incubated with 5 μg of anti-m6A antibodies (Synaptic Systems) previously bound to A/G protein agarose beads (SantaCruz) in IP buffer (10 mM Tris-HCl pH 7.4, 150 mM NaCl, 0.1% Igepal CA-630), and supplemented with RiboLock™ (Thermo Scientific), during 6 hours at 4°C. Beads were washed three times with IP buffer, meRNAs were eluted in elution buffer (10 mM Tris-HCl pH 7.4, 150 mM NaCl, 0.1% Igepal CA-630, 6.67mM m6A) and purified through RNeasy columns (Qiagen) following the manufacturers protocol. For qPCR analyses, (target-meRIP/spike-meRIP)/(target-input/spike-input), were represented per each region. Libraries for massive sequencing were generated using TruSeq Stranded Total RNA Library Prep (Illumina), without previous rRNA depletion, and sequenced by 1×75 single reads at the Centre for Genomic Regulation.

### m6A quantification by dot-blot

5 μg of cheRNA or total RNA for each condition were denatured at 95°C for 5 min and transferred to a Hybond-XL membrane (Amersham) in a Bio-Dot® Microfiltration Apparatus (Bio-Rad) following manufacturer’s intructions. The RNA was UV-crosslinked to the membrane in a Stratalinker® 1800 (Stratagene) at 120 mJoule/cm2. The signal detection was performed after hybridation with anti-m6A antibody (Synaptic Systems), using standard ECL detection reagents.

### CheRNA-seq analysis

Reads were aligned to the mm10 reference genome and to the Luciferase coding sequence using Tophat2 (60) with standard parameters. bedGraph files loaded in the IGV browser were generated with the Bedtool genomecov. The scores of these files were normalized with the total number of aligned reads for each experiment. For the transcriptome assembly, reads coming from six experiments (three WT and three H1-TKO replicates) were pulled, and separated in two files depending on the template strand (Watson or Crick), discriminating them with Samtool view -F 0×10 or -f 0×10, respectively. Spliced reads were discarded from the pull, by removing the entries with a CIGAR string which contained any ‘N’ character. The remaining reads were used to assemble a “genome-guided” transcriptome with Cufflinks v2.2.1 (66). This transcriptome was further curated with home-made scripts to remove low abundance transcripts (minimum coverage < 2.5), remove very short transcripts (size < 300 bp), merge proximal transcripts in the same strand (distance < 2.5 kb) and split transcripts which included an already annotated TSS in the RefGene database. These transcripts were classified in four groups: coding, PROMPTs, lncRNAs and internal antisense transcripts (IASs), according to the diagram shown in **Supplementary Figure 1b**. After transcriptome assembly, 21702 coding transcripts, 3139 PROMPTs, 12673 lncRNAs and 2904 IASs were detected. For the differential gene expression analysis, the quantification of reads per transcript was performed with Salmon (64) using standard parameters. To select differentially expressed genes, DESeq2 software (65) was used, setting two different thresholds: adjusted p-value < 0.01 and fold-change > 2. GO-term enrichment analyses were performed using Panther v14.1 software (66). To account for transcription factor and epigenetic enrichments, Enrichr software was employed (67).

### Histone H1 knock-down RNA-seq analysis

Reads from total RNA-seq preparations (19) were aligned to the hg19 reference genome using Tophat2 with standard parameters. For the transcriptome assembly, reads coming from six experiments (two controls, two H1.4-KD and two multi-KD) were pulled, and separated in two files depending on the template strand (Watson or Crick), discriminating them with Samtool view -F 0×10 or -f 0×10, respectively. The reads that matched a RefGene annotated coding gene were removed from the pull, and the remaining reads were used to assemble a “genome-guided” transcriptome with Cufflinks v2.2.1, exclusively from the noncoding part of the genome. This transcriptome was further curated with homemade scripts to remove low abundance transcripts (minimum coverage < 2.5), remove very short transcripts (size < 300 bp), merge proximal transcripts in the same strand (distance < 2.5 kb) and split transcripts which included an already annotated TSS in the RefGene database. Finally, it was merged with the ENSEMBL coding transcriptome using Cuffmerge, and the transcripts were classified in the four same types as before: in the end, 22827 coding transcripts, 2420 PROMPTs, 14843 lncRNAs and 4562 IASs were detected. The differential gene expression analysis was performed as described above. In this case, the statistical thresholds were set as adjusted p-value < 0.1 and fold-change > 2.

### RNAP II ChIP-seq analysis

Reads were aligned to mouse mm10 and human hg19 reference genomes using bwa mem algorithm. In addition to the standard total read number normalization, the ratio between mouse and human reads was used to correct the H1-TKO cells metaplot signal, according to this formula:

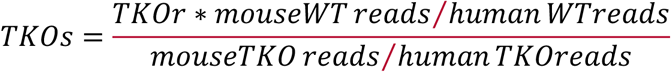

where

TKO_s_ is the spike-in normalized RNApolII signal

TKO_r_ is the total reads normalized RNApolII signal

### ChMeRIP-seq analysis

Reads were aligned to the mm10 reference genome using Tophat2 (60) with standard parameters to generate the bedGraph files loaded in the IGV browser. For the quantification of m6A methylation, the number of reads per transcript was quantified, both in the meRIP and the cheRNA input, using Salmon (64). The methylation level of each transcript was defined as the ratio of the RPKMs in meRIP divided by the RPKMs in the input.

## SUPPLEMENTAL INFORMATION

Supplemental Information including 6 Supplementary Figures and 1 Table can be found with this article online at

## ACKNOWLEDGEMENTS

We are indebted to Arthur Skoultchi and Encarna Martínez-Salas for reagents and advice, Ana Losada and Ana Cuadrado for help with RNAPII-ChIP, Luciana Gómez-Acuña for help on TT-seq, Sandra Benavente for advice in m6A-IP and FTO inhibition, and César Cobaleda for support on computational analysis. We are grateful to the SMOC and Genomic Services at CBMSO and Pepe Belio for art work. We also thank Wendy Bickmore, Andrew Wood and Andrew Jackson for continuous support during MG sabbatical stay at the HGU, and Ferran Azorín, Jordi Bernues, Victor Corces, Crisanto Gutierrez and members of MG lab for critical reading of the manuscript. Work at MG lab was supported by the Spanish Ministry of Sciences and Innovation (BFU2016-78849-P and PID2019-105949GB-I00, co-financed by the European Union FEDER funds), a CSIC grant (2019AEP004), and a Salvador de Madariaga mobility grant (PRX19/00293). JMF-J and CS-M were supported by the Spanish Ministry of Sciences and Innovation fellowships (BES-2014-070050 and BES-2017-079897, respectively) and MS-P was supported by an AGAUR-FI predoctoral fellowship co-financed by Generalitat de Catalunya and the European Social Fund. AJ was supported by the Spanish Ministry of Sciences and Innovation (BFU2017-82805-C2-1-P and PID2020-112783GB-C21) and JFC by Core funding to the MRC Human Genetics Unit from the Medical Research Council (UK).

## AUTHOR CONTRIBUTIONS

JMF-J, CS-M, AF-L and MS-P performed experiments. JMF-J performed all computational analyses. JMF-J, CS-M and MG designed and analized the experiments. MMM contributed to the TT-seq experimental design and analysis. MG conceived the project, analyzed experiments and wrote the article. MG, AJ and JFC secured the funding. All authors analyzed the data, discussed the results and approved the final version of the manuscript.

## LEAD CONTACT AND MATERIALS AVAILABILITY

Further information and requests for resources and reagents should be directed to and will be fulfilled by the Lead Contact, María Gómez

