## Supplementary figures and images for "Histone H1 regulates non-coding RNA turnover on chromatin in a m6A-dependent manner"

### Supplementary Figure 1

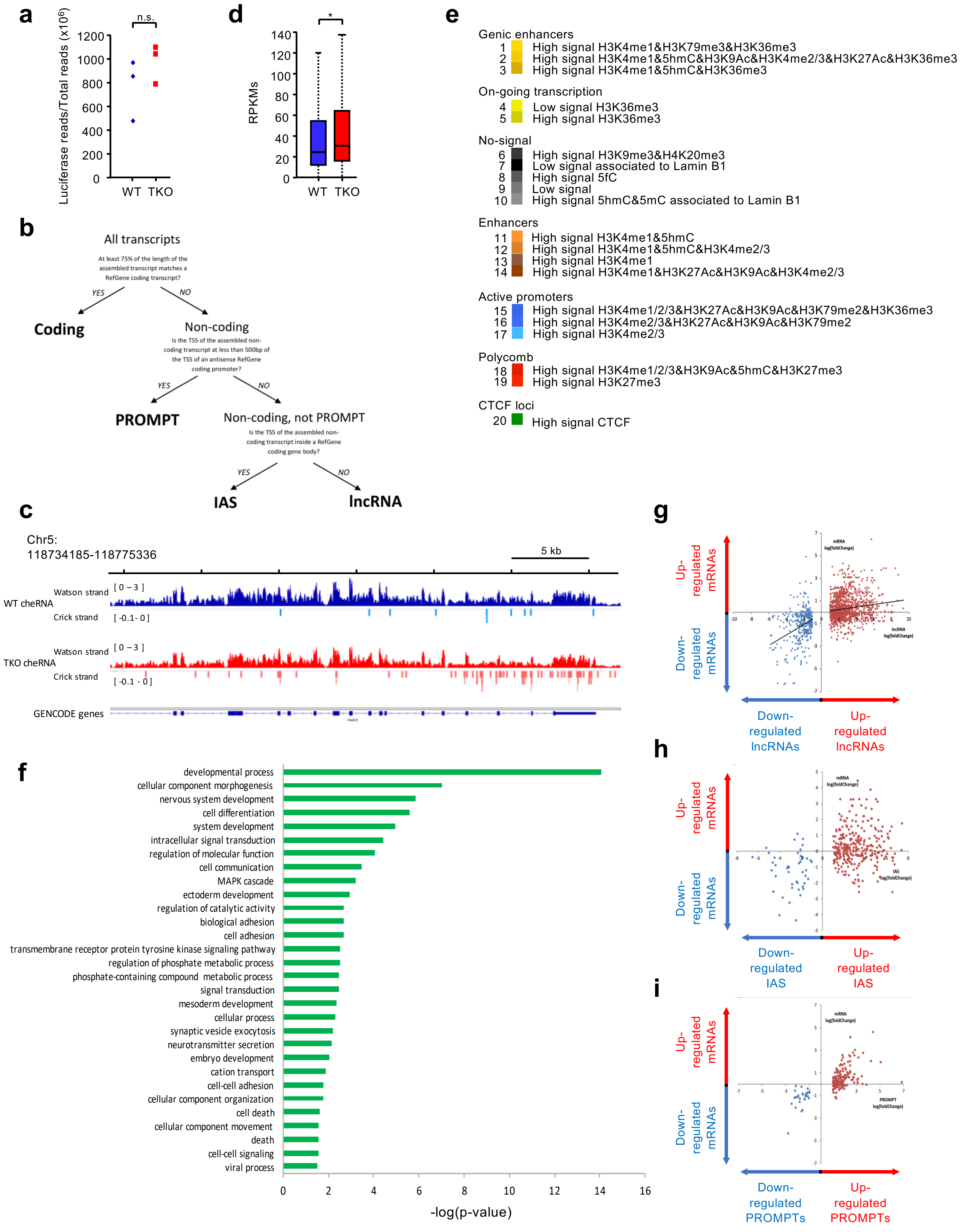


**Supplementary Figure 1**

### Supplementary Figure 2

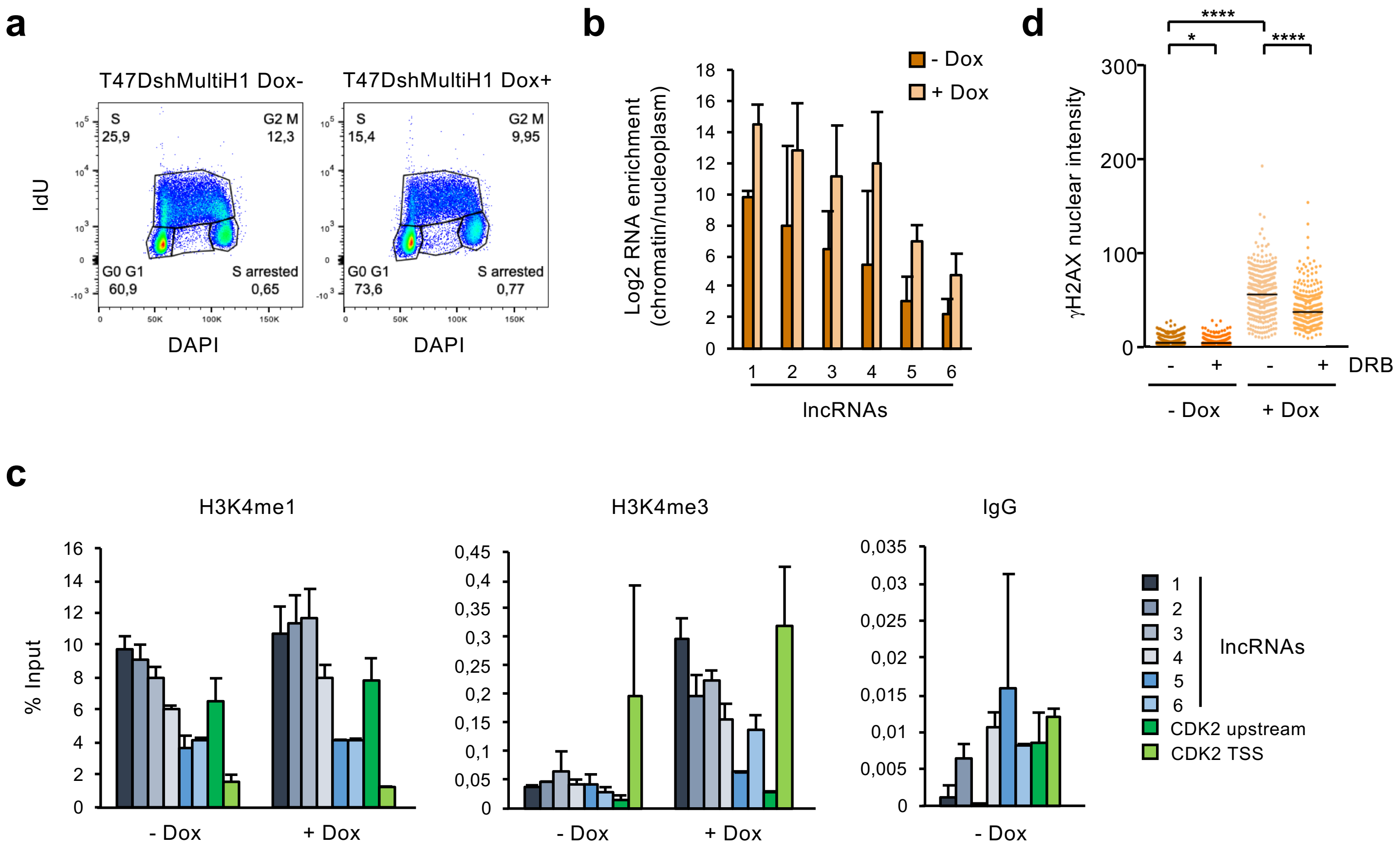


Supplementary Figure 2

### Supplementary Figure 3

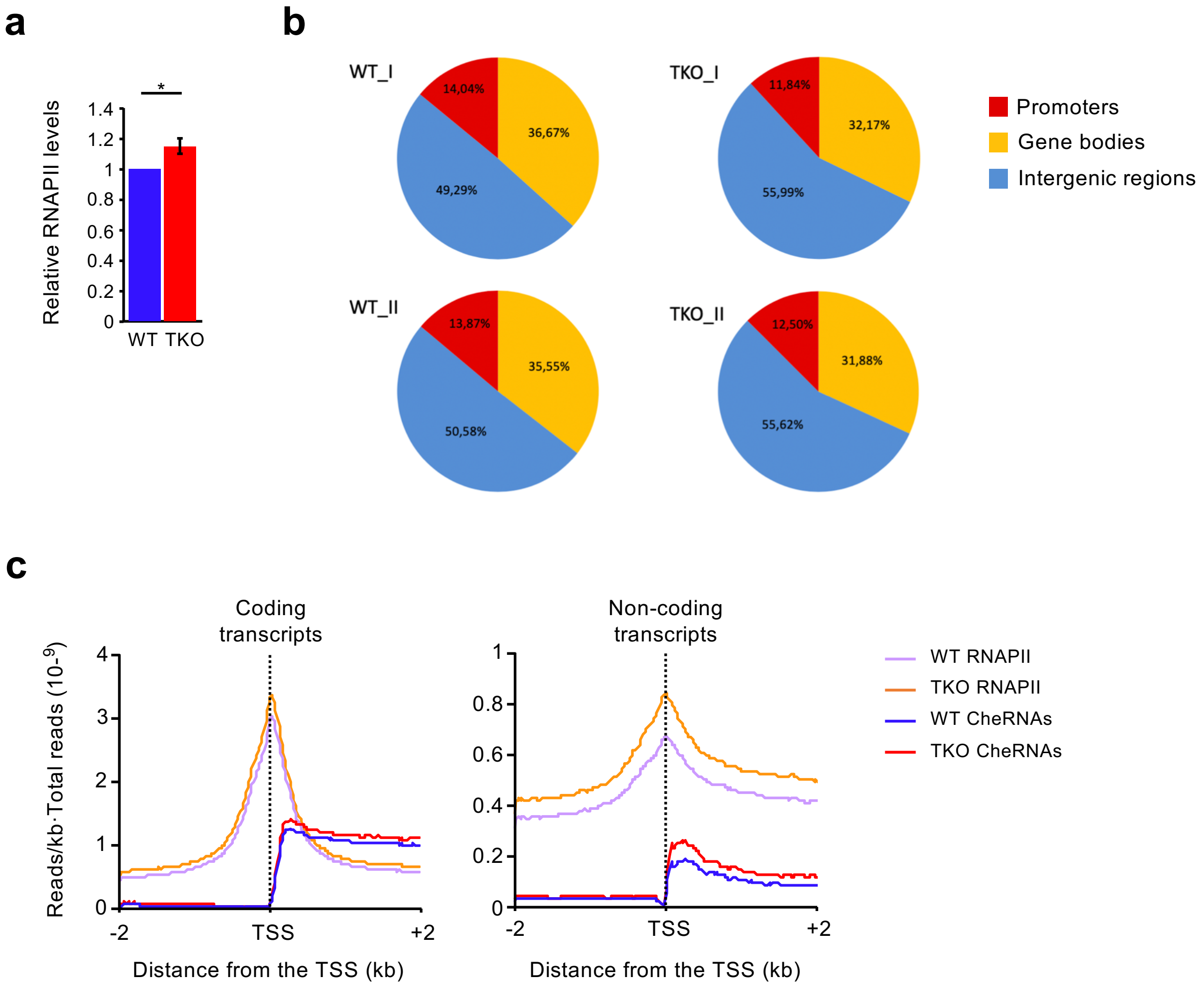


**Supplementary Figure 3**

### Supplementary Figure 4

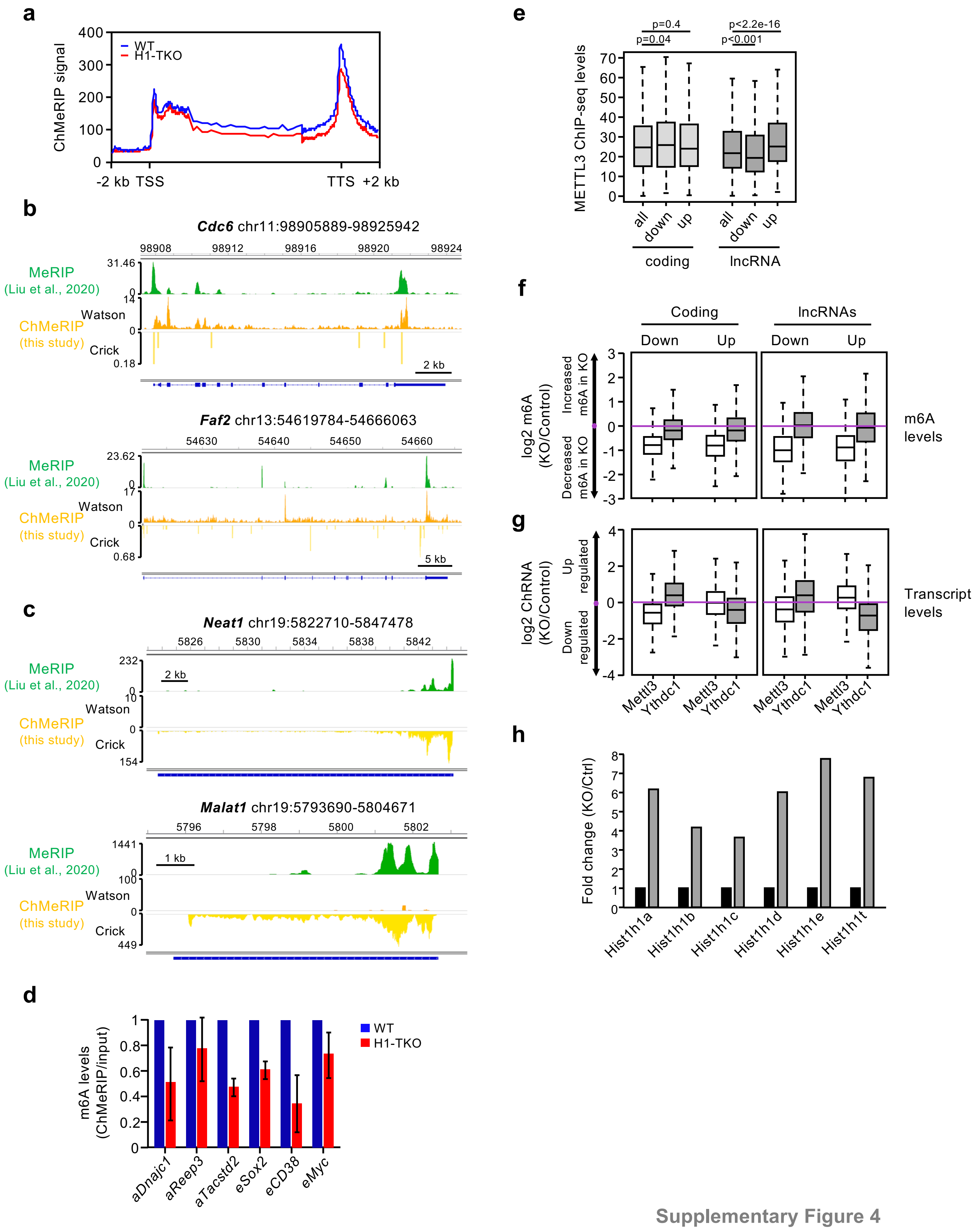


Supplementary Figure 4

### Supplementary Figure 5

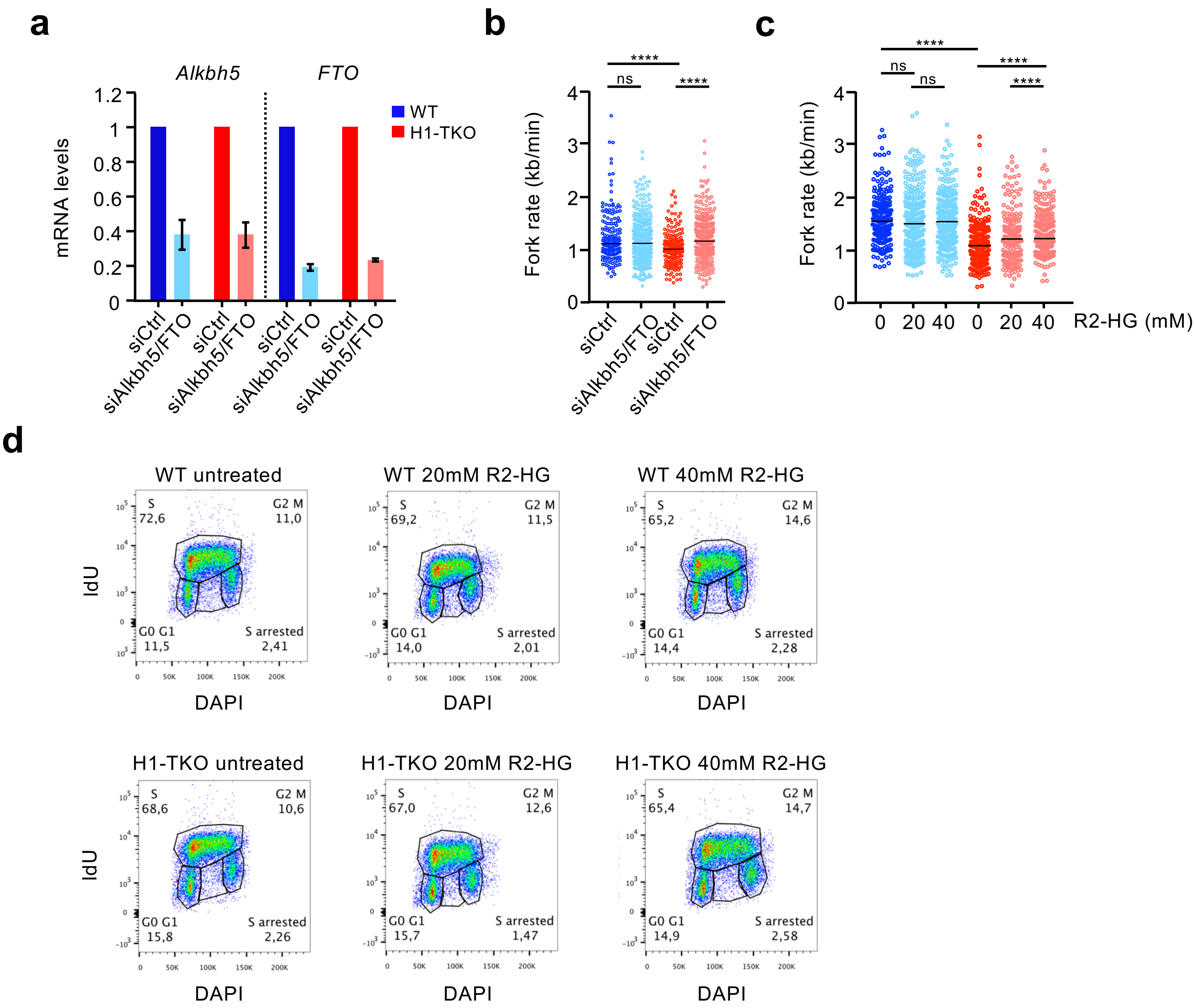


**Supplementary Figure 5**

### Supplementary Figure 6

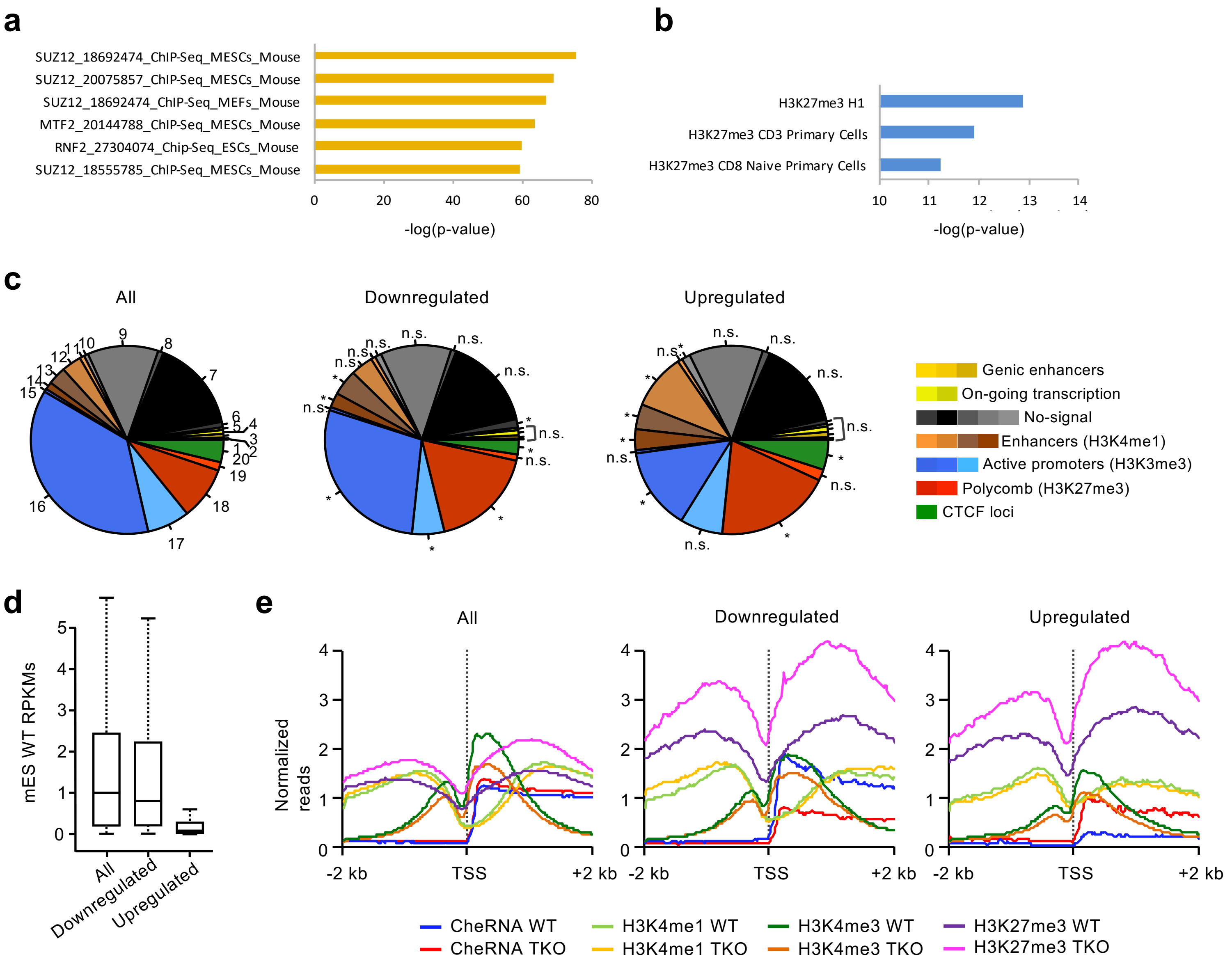


**Supplementary Figure 6**
