## Supplementary Figure Legends for "Histone H1 regulates non-coding RNA turnover on chromatin in a m6A-dependent manner"

**Supplementary Figure 1. Differential CheRNA-seq analyses between WT and H1-TKO mESCs reveals ncRNA enrichment in H1-depleted chromatin. (a)** Percentage of Luciferase reads in WT and H1-TKO CheRNA-seq replicates. n.s., not significant, t-test p-value=0.30. **(b)** Diagram of transcript classification after the "genome-guided" transcriptome assembly (see Methods for details). A total number of 21,702 mRNAs, 3,139 PROMPTs, 12,673 lncRNAs, and 2,904 IAS were detected. **(c)** Representative IGV browser snapshot of a region not assembled as a transcript with differential CheRNA reads accumulation in H1-TKO cells in the Crick strand. **(d)** Boxplot showing the distribution of RPKMs at inter-transcript regions in WT and H1-TKO cells *Mann-Whitney-Wilcoxon test p-value<$2.2x10^{-16}$. **(e)** Key for mESCs chromatin states as in Juan et al. (2016). **(f)** Panther GO-term enrichment analysis of differentially accumulated coding transcripts. **(g)** Fold-change of lncRNA expression between WT and H1-TKO cells (X-axis) and the nearest coding gene (Y-axis). **(h, i)** Same as in (g) for PROMPTS (h) or IAS (i).

**Supplementary Figure 2. lncRNAs are overexpressed upon inducible knock-down of histone H1 genes in human cells. (a)** Representative cell cycle distribution of T47D shmultiH1 cells without (-Dox) and with 3 days of doxyclyclin treatment (+Dox). The percentage of actively replicating cells was evaluated by 20 min pulse with IdU. Upon 3 days knock-down of histone H1 genes T47D shmultiH1 cells partially accumulate in G1 as described (Izquierdo-Boulstridge et al., 2017). (**b**) Chromatin/nucleoplasm enrichment of 6 lncRNAs in T47D shmultiH1 -/+ Dox. RT-qPCR data represent mean and standard deviation of four independent RNA preparations, expressed as ratio relative to nucleoplasm (-Dox). (n=4) (**c**) ChIP-qPCR abundances of H3K4me1 and H3K4me3 histone marks at the indicated lncRNAs upon Dox treatment. A representative experiment is shown. IgG was used as ChIP non-specific control. Primer pairs for CDK2 TSS and 3kb-upstream were used as enrichment controls for H3K4me3 immunoprecipitation. (**d**) Distribution of γH2AX nuclear intensities in T47D shmultiH1 Dox- and Dox+ cells treated (+) or not (-) with DRB for 3h, n > 311.

**Supplementary Figure 3. Accumulated transcripts are tethered to chromatin by RNAPII. (a)** Normalized amounts of immunoprecipitated RNAPII molecules in WT and H1-TKO cells in both ChIP-seq replicates. *z-test p-value=$1.6 \cdot 10^{-6}$. **(b)** Pie plots showing the distribution of RNAPII reads in different genomic regions in both ChIP-seq replicate experiments. Promoter annotation was downloaded from GENCODE coding transcripts database: the locus comprises TSS +/- 1kb, gene bodies encompass the region from TSS+1kb to TTS, and intergenic regions are the remaining genome. **(c)** Metaplots of CheRNA-seq and RNAPII-seq spike-in normalized signals from duplicate experiments in WT and H1-TKO cells, plotted in a 4kb window around the TSS. Note that the scale of the Y-axis is not the same in both plots, in order to facilitate the visualization of RNAPII signal differences between WT and H1-TKO conditions. RNAPII WT and TKO signals were multiplied by a scale factor of 1:2, to facilitate the visualization in a single plot.

**Supplementary Figure 4. m6A alterations of chromatin-enriched transcripts in H1 TKO cells. (a)** Metagene profiles of ChMeRIP reads along coding transcripts in two replicates for WT and H1-TKO mESCs. **(b, c)** IGV browser snapshots of MeRIP (Liu et al., 2020) and ChMeRIP normalized reads along the genes *Cdc6* and *Faf2* (b), and the lncRNAs *Neat1* and *Malat1* (c). **(d)** m6A levels of 6 lncRNAs quantified through normalizing ChMeRIP qPCR results with spike-in between WT and H1-TKO cells. n=3 biological replicates; error bars indicate means +/- SEM. **(e)** METTL3 levels at a 4kb window surrounding the TSS of the indicated transcript categories. METTL3 ChIP seq data are from Xu et al (2021). Statistical differences between distributions were assessed with the Mann-Whitney rank sum test. P values are indicated. **(f)** Plots showing the ratios of m6A content between Mettl3-KO (white) or Ythdc1-KO (grey) mES cells and their WT counterparts at the indicated transcript categories. **(g)** Same as (f) for RNA levels. m6A RIP-seq and RNA-seq data are from Liu et al (2020). In all cases boxplots denote the medians and the interquantile ranges and the whiskers represent the 10 and 90 percentiles. **(h)** Histogram plot showing the fold-change expression of histone H1 variants in *Mettl3*-KO mES cells. RNA-seq data are from Liu et al. (2020).

**Supplementary Figure 5. Inhibiting m6A demethylase activity rescues the slow replication phenotype of H1-TKO cells. (a)** RT-qPCR mRNA silencing levels of *Alkbh5* and *Fto* 24h after cellular transfection with siCtrl or a cocktail of siAlkbh5/Fto. mRNA levels were normalized to *Hprt* mRNA levels at each condition. Data show the median +/- s.d of two independent replicates (n=2). **(b, c)** Replication fork rates of WT and H1-TKO cells transfected with the indicated siRNAs (b), or treated with the indicated amounts of R2-HG for 24h (c). Median values are indicated. Data shown are pooled from two independent experiments (n > 350). Statistical differences between distributions were assessed with the Mann-Whitney rank sum test. p-value: ****<0.0001. (**d**) Representative cell cycle distribution of WT or H1-TKO mES cells treated for 24h in the presence of the indicated amounts of R2-HG. The percentage of actively replicating cells was evaluated by 20 min pulse with IdU.

**Supplementary Figure 6. Characterization of differentially accumulated coding transcripts. (a, b)** Enrichr analysis of overrepresented transcription factor binding sites (**a**), or chromatin marks (**b**), at the promoter regions of differential coding transcripts. The -log(p-value) for each ChIP-seq database entry is plotted. **(c)** Pie plots showing the percentage of transcripts whose promoters matches each chromatin state. The percentage was compared with the expected one obtained from 100 random permutations of the differential transcripts, and the p-value was calculated. *p-value<0.01; ns, not significant.

Chromatin states are from Juan et al. (2016). **(d)** Boxplots showing the distribution of the RPKMs of differential coding transcripts in WT mES cells. **(e)** Metaplots of the indicated ChIP-seq signals and CheRNA-seq reads for WT and H1-TKO cells, plotted in 4kb windows around the TSS. H3H4me1 WT and H1-TKO signals were multiplied by a scale factor of 2, to facilitate the visualization in a single plot. ChIP data are from Geeven et al (2015).
